# Marine protection enhances the resilience of biological communities on temperate rocky reefs

**DOI:** 10.1101/2023.10.26.564208

**Authors:** Jose A. Sanabria-Fernandez, Josu G. Alday

## Abstract

Conservation science faces the urgent challenge of halting the biodiversity loss caused by the biological crisis of the present era. To achieve this, conservation science requires cutting-edge tools to focus on vital properties of ecosystems, such as the resilience. Resilience informs about the cost of recovering biological communities. Here, we developed a metric to quantify the communities ecological recovery cost based on the multidimensional distance between unprotected and partially protected communities from totally protected communities in Cabo de Gata Marine Reserve. Our results show that the biological community composed of fish, macroinvertebrates and cryptic fish, and macroalgae species in unprotected zones requires a higher ecological recovery cost than in partially protected zones when moving towards a fully protected community. This research contributes to monitoring marine the effectiveness of marine protection from a resilience perspective, with the goal of promoting the use of the recovery cost metric for building resilient coastal ecosystems.

## 1. Introduction

Biodiversity conservation is one of the principal needs of our times since human activities are causing swift, profound, and ongoing environmental changes that affect the earth’s system (Scholes 2016). These abrupt changes are leading the earth to an alternate state with adverse consequences for humanity (Scheffer et al., 2001; Lenton et al., 2008; Barnosky et al., 2011), making an undoubted argument for the beginning of a new era, the Anthropocene (Crutzen and Stoermer, 2000; Steffen et al., 2011; Lewis and Maslin, 2015). Out of nine planetary boundaries defined to prevent a global phase shift, biodiversity loss has surpassed its safety threshold (Rockström et al., 2009). Current extinction rates are 100 to 1000 times larger than background extinction rates (Alroy, 2015), providing support to the sixth global mass extinction (Barnosky et al., 2011; Dirzo et al., 2014; McCauley et al., 2015). Since the evolution of new species usually takes thousands of generations (Gavrilets, 2003), we may be facing a long-lasting biodiversity loss and an associated decline in ecosystem services. Thus, there is an urgent need to implement effective conservation measures to halt biodiversity loss, safeguard our best natural resources, and restore biodiversity and ecosystem services (IPBES, 2019; EU2030 Biodiversity Strategy, 2020). Not surprisingly, biodiversity conservation goes beyond the academic box and has become an integral part of society with a presence in (most) political agendas (United Nations 2000; Youatt, 2015; Lobo and Jacques, 2017).

Fueled by the Convention on Biological Diversity (CBD), the designation of protected areas has emerged as a relevant management tool to counteract current biodiversity trends (Secretariat of the CBD, 2014). The traditional nomination of protected areas combines multiple biological, social, and aesthetic criteria to target biologically valuable areas that maximize conservation benefits (Roberts et al. 2003; Selig et al., 2014; Mcleod et al., 2019). Protected areas, terrestrial and marine (hereafter, MPAs), are designed to preserve biodiversity (in a broad sense), enhance species biomass, and increase the recovery of threatened fished species (Halpern, 2003; Edgar et al., 2014). But beyond these specific benefits, the MPAs also could strengthen vital community properties, such as resilience, to maintain the ecosystem healthy (Mellin et al., 2016). In fact, resilience plays an essential role in the health of natural ecosystems, as it could prevent the occurrence of catastrophic and abrupt phase changes in natural ecosystems (Gunderson et al., 2009; Scheffer, 2009). However, despite its relevance, the designation of protected areas often fails to consider their resilience (Game et al., 2008; Abelson et al., 2016). Therefore, considering resilience in the MPAs implementation would contribute to improving and keeping them healthier (Green et al., 2009).

Resilience is the capacity of a system to absorb disturbances and reorganize while the ongoing change retains essentially the same function, structure, identity, and feedback (Walker et al., 2004). This definition is multidimensional because it integrates persistence, abundance, resistance, and the existence of local asymptotic stability at multiple equilibria (Donohue et al., 2016). Therefore, resilience is a multi-faceted concept that includes multiple components. From a theoretical perspective, resilience is lost when a healthy community is damaged and approaches a degraded state (Folke et al., 2004). Conceptually measuring the distance between healthy and degraded communities could inform us about the space gap between both communities. Indeed, we could estimate the ecological recovery cost using the multidimensional distance from degraded (i.e., unprotected) to healthy communities (i.e., fully protected). Here, we understood the ecological community recovery cost based on changes in the presence and density of species, measured as the multidimensional distance between unprotected and fully protected communities. For example, two biological communities (unprotected and fully protected) separated by a small distance imply low ecological recovery cost due to their similar community composition. However, if the multidimensional distance between communities is high, it would suggest a high ecological recovery cost, resulting from marked differences in the composition of both communities. Thus, the development of metrics that shed light on the dimensions of resilience could have a significant impact on the proper functioning of protected areas. Unfortunately, it is a scarcely studied topic (Maynard et al., 2010; Maynard et al., 2015), but profoundly important for safeguarding biodiversity in the current biological crisis era, since this would help us learn about the recovery cost of biodiversity before enforcing conservation measures.

In this context lies the need for our study, which aims to report the application of a novel metric based on the ecological recovery cost of biological communities. Additionally, it seeks to investigate the role that a lack of protection from anthropogenic activities plays in the recovery costs of marine communities. To calibrate this approach, we studied the marine biodiversity in fully, partially, and unprotected zones of the Cabo de Gata Marine Reserve and its adjacent waters. First, we characterized the biodiversity composition of four community categories, i.e., fish, macroinvertebrates and cryptic fish, and macroalgae communities, and the overall community composition (combination of the three of them). Second, we quantified the ecological recovery cost required to transform the partially protected and unprotected zones into a totally protected zone. Third, we investigated the species responsible for shifts in recovery costs, i.e., species whose presence or density depends on the protection status. This approach could be used in monitoring and designing terms in the field of conservation. For instance, as we presented here, it allows for revealing the well-functioning of the protected zones. But also, this metric has a potential application in the design of a new protected area since this approach could unveil priority geographic areas to preserve. Therefore, the metric presented could assess and guide the monitoring process of the protected areas from the perspective of resilience, helping to achieve conservation goals and increasing the efficiency of protective measures.

## 2. Methods

### 2.1. Study area

We sampled 28 locations situated in Cabo de Gata marine reserve and its adjacent unprotected area, located in the southeast of the Iberian Peninsula. We surveyed eight locations in the no-take zone (the fully protected zone), ten locations in the partially protected area, and ten locations in unprotected areas adjacent to the marine reserve (Fig. 1). In this article, fully protected zones (hereafter FPZs) refer to areas with severe restrictions and levels of surveillance within the marine reserve (no-take zones). In FPZs, all anthropogenic extractive and recreational activities are entirely prohibited, indicating that these areas maintain a healthy state of marine biodiversity (Edgar et al., 2014, Sala and Giakoumi, 2017). Therefore, in the present study, we selected FPZs as reference zones for assessing the ecological community recovery cost. Partially protected zones (hereafter PPZs) refer to areas within the marine reserve where certain anthropogenic activities, such as recreational fishing and diving, are permitted. Finally, unprotected zones (UNZs) lie outside marine protected areas and are referred to as areas that lack surveillance, and anthropogenic activities are allowed. During the summer season of 2014, we conducted underwater surveys on rocky reefs ranging from 6 to 10 meters in depth.

**Figure 1.**
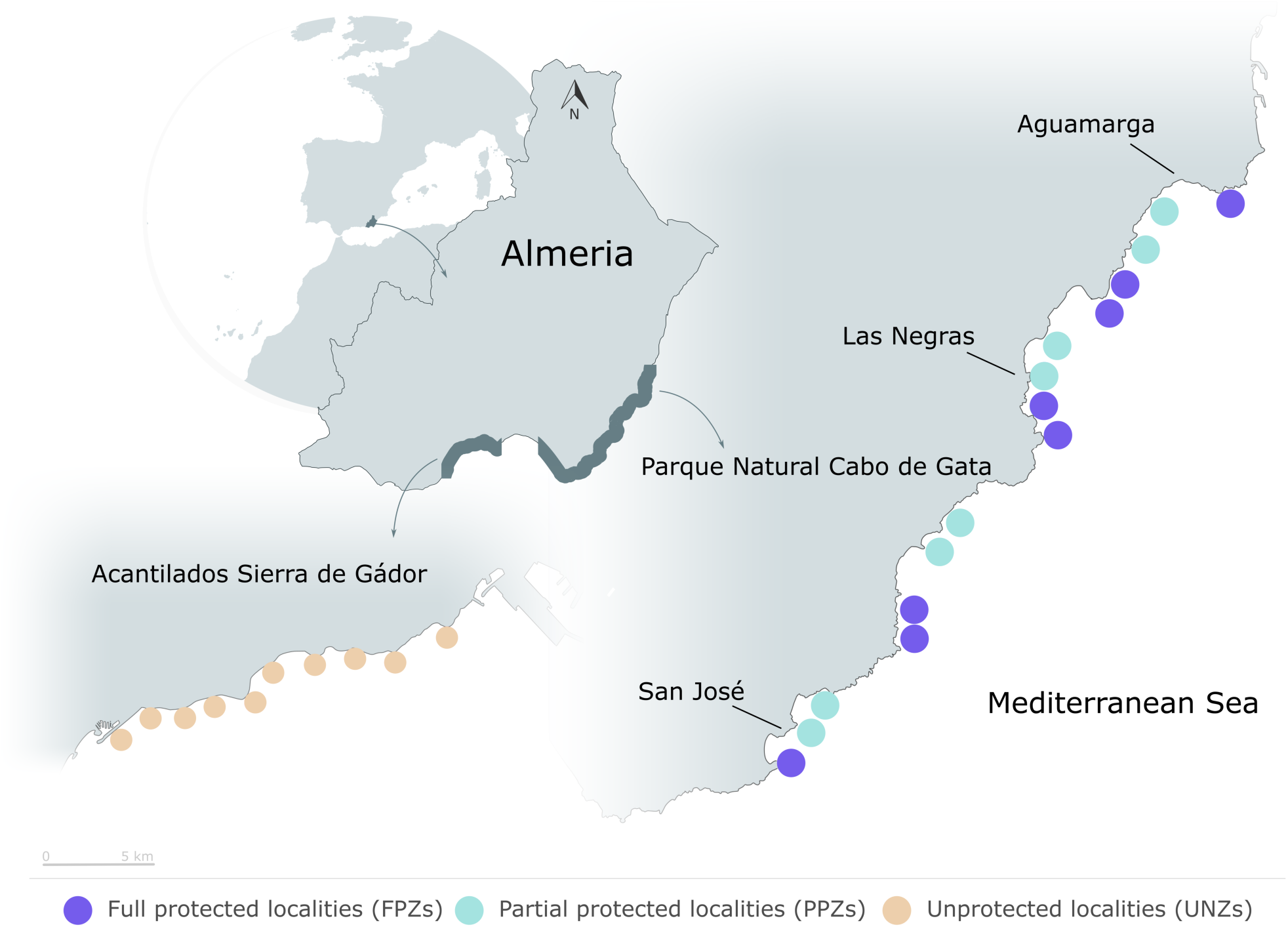
Map of the study area in Cabo de Gata marine reserve and adjacent waters, located in the Mediterranean Sea. Circles show the sampled zones (N=28), colors stand for their protection status.

### 2.2. Data collection

We collected the reef associated marine biodiversity data using the Reef Life Survey standardized underwater visual census protocol (Edgar and Stuart-Smith, 2014). We selected this protocol because it is a worldwide standardized protocol, it is a non-destructive and non-extractive methodology, so we did not remove any animals or algae during sampling. Reef Life survey methodology allows us to obtain information on fish, macroinvertebrates and cryptic fish, and macroalgae in a single scuba dive. The method involved a 50 m belt transect over which we applied three different approaches (Supplementary Fig.1). In the first approach, we focused on the fish community, quantifying the species density in a total area of 500 m^2^ (50×10-m^2^). In the second approach, we counted the macroinvertebrates and cryptic fish species and their density within 100 m^2^ (50×2-m^2^). In the third approach, we took 20 high-quality pictures of the seabed spread along the 50 m transect and separated by 2.5 m to quantify the coverage of macroalgae (Supplementary Fig.1). In each picture, we draw a centered digital square of 400 cm^2^ to randomly select 20 points (Sanabria-Fernandez et al., 2018) using CPCe software (Kohler and Gill, 2006). After classifying the macroalgae on which the random points fell, we calculated their percent cover as the number of species points over the total number of points for each transect. In this way, we obtained three subsets of data for each sampled location, i.e., fish, macroinvertebrates and cryptic fish, and macroalgae.

In the context of the abiotic dimension, we selected a subset of the 14 most important variables that condition the marine biological communities in temperate rocky reefs (Tittensor et al., 2010, Stuart-Smith et al., 2013). In detail, we used the location coordinates to collect the scores of the variables from the Bio-Oracle database (Assis et al., 2018) and (Yeager et al., 2017) using the extract function of the raster R package. We collected the minimum, mean, and maximum of Sea Surface Temperature (°C), salinity mean (PSS), pH, nitrate concentration (mmol·m−3), silicate concentration (mmol·m−3), phosphate concentration (mmol·m−3), as well as minimum, mean, and maximum of chlorophyll concentration (mg m−3), photosynthetically available radiation (Einstein/m2/day), the wave energy, all of them with a resolution of 9.2 × 9.2 km, and the depth of the localities.

### 2.3. Data analyses

Previous to conducting the analyses, we performed a Hellinger’s transformation on the fish and macroinvertebrates and cryptic fish community to reduce the influence of high-density species (Oksanen et al., 2022). We also log-transformed (x+1) the percent cover of macroalgae (Oksanen et al., 2022). Subsequently, we elaborated a dissimilarity matrix based on the Bray-Curtis distances for each community, i.e., fish, macroinvertebrates and cryptic fish, and macroalgae communities. Finally, we combined the three distance matrices (containing data with various methods and units) to obtain a single community dissimilarity matrix using the fuse function from the analogue R package. Then, we used the species richness, i.e., the number of species of each taxonomic group (fish, macroinvertebrates and cryptic fish, and macroalgae) to compute a weighted sum of the dissimilarity matrices (Bennion et al., 2015). Thus, the subsequent analyses of the overall community composition relied on this fused species matrix.

To achieve the study goal, we conducted four sets of analyses. Firstly, we investigated whether the geographic distance and environmental variables could be factors that significantly modulate the composition of biological communities in the study area. Initially, we explored the correlation of the geographic position in order to see how communities are far from each other in geographic space, fitting a Mantel test for each biological dissimilarity matrices, i.e., fish, macroinvertebrates and cryptic fish, and macroalgae, and overall community *versus* the Euclidean distance matrix of the location coordinates. To do that, we applied the mantel function of the vegan package, using Pearson’s correlation coefficient and 999 permutations. After that, we studied whether the abiotic dimension composed by 14 variables conditions the biological communities at our study scale, while controlling the effect of the geographic position to remove any possible spatial autocorrelation. To do so, we fitted a partial mantel function for each biological matrix, using the partial.mantel function from the vegan package, applying Pearson’s correlation coefficient and 999 permutations.

Secondly, we explored the effect of the protection levels (FPZs, PPZs, and UNZs) on the biological community composition, i.e., fish, macroinvertebrates and cryptic fish, and macroalgae, and overall community composition. To do so, we ran a Permutational Multivariate Analysis of Variance (PERMANOVA) using the adonis function from vegan package, on each community matrix separately (fish, macroinvertebrates and cryptic fish, macroalgae), and on the fused species matrix (overall community), while the following pair-wise comparisons between protection status were Bonferroni corrected (Sokal and Rohlf, 1995).

In the third set of analyses we focused on quantifying the ecological community recovery cost of the four biological communities (fish, macroinvertebrates and cryptic fish, macroalgae, and overall community composition) between the three different protection levels. Here, we define ecological recovery cost as the multidimensional distance from the UNZs and PPZs zones to the confidence interval (CI 95%) of the FPZ centroid (Fig. 2, Anderson, 2006). To do so, we calculate the distances from each location to the confidence interval of the different community centroids in the 3-dimensional ordination space. This can be defined as a multivariate distance-based metric that measures the rate of change in community structure based on the presence or absence of species and their density. This provides a straightforward proxy measure of the ecological community recovery cost scores oscillating between 0 to 1, increasing as it approaches 1. Specifically, we quantified this metric setting the UNZs and PPZs as starting points and the confidence interval (CI 95%) of the FPZ centroid as the arrival point (Fig. 2). We selected the FPZs as reference zones for assessing the ecological community recovery cost because they maintain a healthy state of marine biodiversity (Edgar et al., 2014, Sala and Giakoumi, 2017) since all anthropogenic extractive and recreational activities are entirely prohibited. Hence, using them as reference zones allows us to gain a deeper understanding of how human activities affect the marine biodiversity. We obtained a unique value for each biological community and protection status performing the average of the multidimensional distance between protection zones.

**Figure 2.**
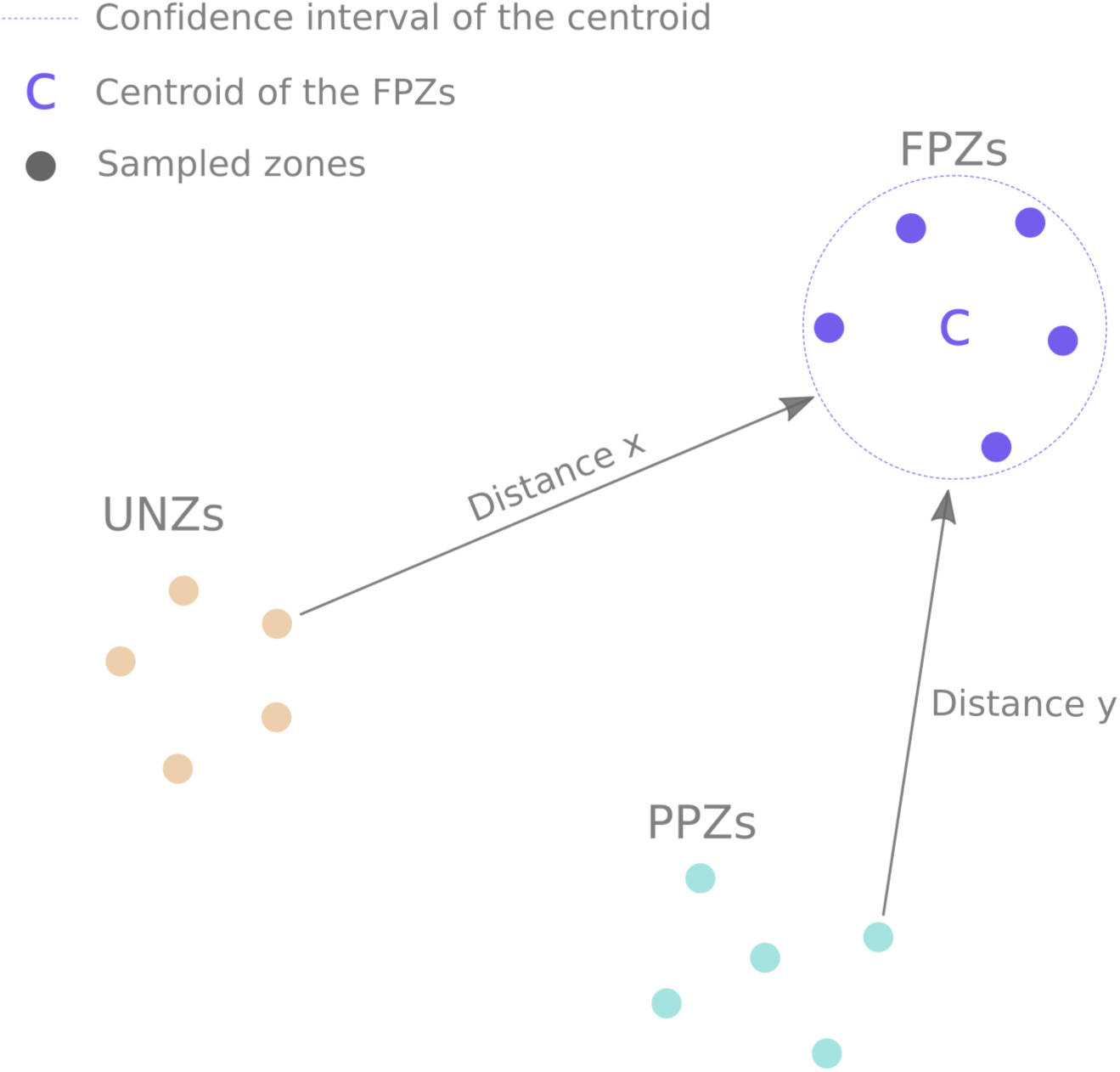
The conceptual framework represents how we measured the multidimensional distances in Euclidean space. For example, the distance “x” represents the space gap between unprotected zones (UNZs) and the confidence interval (CI 95%) of the centroid of fully protected zones (FPZs). Additionally, we depicted the distance “y”, as the spatial gap between partially protected zones (PPZs) and the confidence interval of the centroid of FPZs. We computed this process for all zones under both protection status and performed the average of the distances for UNZs and PPZs.

Lastly, our fourth set of analyses explored the effects of the protection levels (FPZs, PPZs, and UNZs) on species and their densities from a univariate perspective. Specifically, we looked for species changing their density or coverage when comparing FPZs to UNZs and PPZs. To do this, we fitted Linear Mixed Models (LMMs) using the lme() function from the nmle package to each species of fish, macroinvertebrate, cryptic fish, and macroalgae, with density or cover as the response variable and the protection status as the sole fixed factor. Additionally, we included the sampling sites as a random factor to account for spatial pseud-replication.

All data preparation and computations developed in the article would not have been possible without the R open-source software (R core team, 2023), including the following packages: analogue (Simpson and Oksanen, 2021), vegan (Oksanen et al., 2022), nmle (Pinheiro and Bates, 2023), and raster (Hijmans et al., 2023). The recovery cost function and an example of how the function works have been included as supplementary material written in R language.

## 3. Results

### 3.1. Effects of the geographic distance and the abiotic variables on the biological community

Considering the fish, macroinvertebrates and cryptic fish, macroalgae, and overall community, the geographical distance did not affect the dissimilarity of biological communities (See Table S1). Equally, there were no significant results in the partial mantel tests correlations between the four biological matrices and environmental variables (See Supplementary Table 1).

### 3.2. Biological community composition under different protection levels

The overall community composition depends on the protection status (permanova, p<0.001, Fig. 3a, Table 1a) and accounted for 35% of the variance in the overall data. So it is for the fish and macroalgae communities, which depend on the protection status in all comparisons, explaining 17% and 31% of the variance respectively (Fig. 3b; Table 1b for fish, and Fig. 3d and Table 1d for macroalgae). The macroinvertebrate and cryptic fish community also changes with the protection status (Fig. 3c, Table 1c), except for the FPZs *vs.* PPZs comparison (R^2^=0.07, p=0.054).

**Figure 3.**
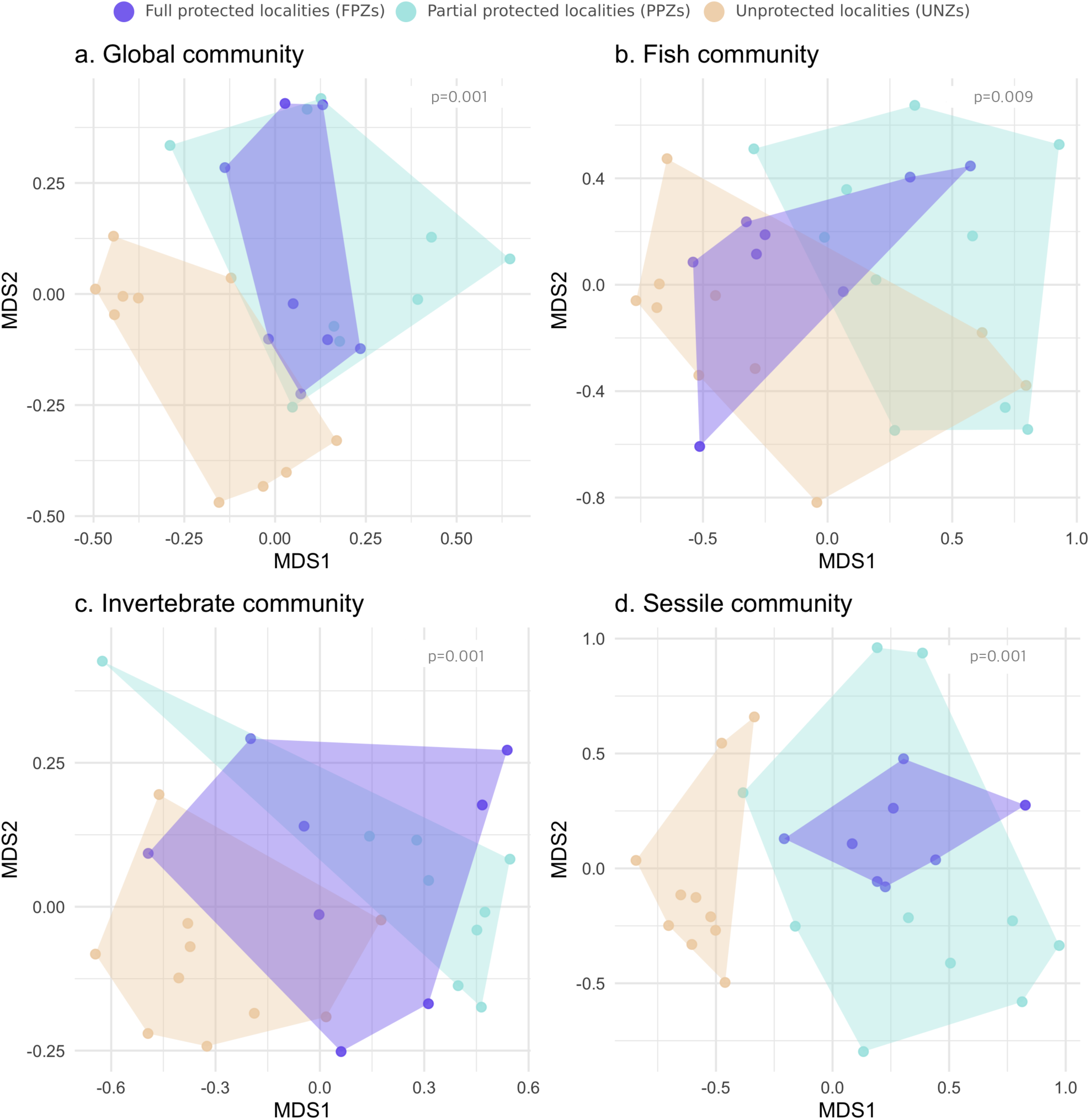
Non-metric multidimensional scaling (NMDS) ordination of species-density matrices from the: a) Overall community, i.e., the combination of fish, macroinvertebrates and cryptic, and macroalgae communities. b) fish community, c) macroinvertebrates and cryptic, and d) macroalgae community in three protection levels in Cabo de Gata. The drawn area represents the convex hull of each protection level.

**Table 1.** Summarized results of Permanova test. Significant differences are marked in bold.

|  | R<sup>2</sup> | F | P |
| --- | --- | --- | --- |
| **a) Overall Community composition** |  |  |  |
| All zones | 0.35 | 6.23 | **0.001** |
| FPZs vs PPZs | 0.11 | 3.90 | **0.006** |
| FPZs vs UNZs | 0.24 | 8.55 | **0.001** |
| PPZs vs UNZs | 0.23 | 8.53 | **0.001** |
| **b) Fish community** |  |  |  |
| All zones | 0.17 | 2.63 | **0.009** |
| FPZs vs PPZs | 0.07 | 2.18 | **0.048** |
| FPZs vs UNZs | 0.10 | 3.08 | **0.013** |
| PPZs vs UNZs | 0.13 | 4.01 | **0.007** |
| **c) Macroinvertebrate and cryptic fish community** |  |  |  |
| All zones | 0.30 | 5.15 | **0.001** |
| FPZs vs PPZs | 0.07 | 2.53 | 0.054 |
| FPZs vs UNZs | 0.23 | 7.77 | **0.001** |
| PPZs vs UNZs | 0.24 | 8.10 | **0.001** |
| **d) Macroalgae community** |  |  |  |
| All zones | 0.31 | 5.52 | **0.001** |
| FPZs vs PPZs | 0.09 | 3.14 | **0.010** |
| FPZs vs UNZs | 0.22 | 7.90 | **0.001** |
| PPZs vs UNZs | 0.19 | 6.20 | **0.001** |

### 3.3. Ecological community recovery cost from UNZs and PPZs to FPZs

As for the ecological recovery cost of the overall community, the UNZs showed higher scores than PPZs when aiming to reach a biological community similar to FPZs (Table 2a). In detail, this recovery cost of the overall community in UNZs was 10% higher than in PPZs. In terms of the fish community, to our surprise, PPZs showed up to 11% higher recovery cost, more than in UNZs (Table 2b). Regarding the macroinvertebrate and cryptic fish community, the ecological recovery cost in UNZs was up to 50% higher than in PPZs. Nevertheless, this recovery cost of PPZs in the macroinvertebrate and cryptic fish community was the lowest found in the study (0.09±0.06). Finally, the macroalgae community reflected the highest ecological recovery cost identified in UNZs (0.52±0.05). This score was the highest recovery cost value found in this study. Summarizing these results, in three of the four comparisons, the recovery cost was higher in UNZs and only once was it higher in PPZs.

**Table 2.** Summarized results of compositional ecological recovery cost. Reported values are average scores ± SE.

|  | Ecological Community<br>Recovery cost |
| --- | --- |
| **a) Overall community** |  |
| PPZs | 0.32±0.07 |
| UNZs | 0.42±0.06 |
| **b) Fish community** |  |
| PPZs | 0.23±0.07 |
| UNZs | 0.12±0.06 |
| **c) Macroinvertebrate and cryptic fish community** |  |
| PPZs | 0.09±0.06 |
| UNZs | 0.18±0.05 |
| **d) Macroalgae community** |  |
| PPZs | 0.44±0.05 |
| UNZs | 0.52±0.05 |

### 3.4. Changes in species composition along protection levels

The changes in ecological community recovery cost between protection levels evidenced fluctuations in the presence, density, or coverage of fish species, macroinvertebrates and cryptic fish, and macroalgae. On the fish community side, species such as *Coris julis*, *Symphodus tinca*, and *Serranus scriba* showed significantly higher densities in FPZs than in UNZs. Similarly, the same pattern was observed with *Chelon auratus*, *Sphyraena viridensis*, and *Mugil cephalus* in FPZs compared to PPZs (Fig. 4a). At the macroinvertebrate and cryptic fish community level, species such as *Paracentrotus lividus*, *Echinaster sepositus*, and *Columbella rustica* evidenced significantly higher densities in FPZs than in UNZs. Likewise, a similar pattern was noted with *Holothuria sanctori*, *Apogon imberbis*, and *Cocinasteria tenuispina* in FPZs compared to PPZs (Fig. 4b). Lastly, at the macroalgae community level, species such as *Halopteris filicina*, *Pseudolithoderma adriaticum*, and *Padina pavonica* showed significantly higher cover in FPZs than in UNZs. Similarly, the same pattern was observed with *Padiva pavonica*, *Hydrolithon farinosum*, and *Feldmannia lebelii* in FPZs compared to PPZs (Fig. 4c).

**Figure 4.**
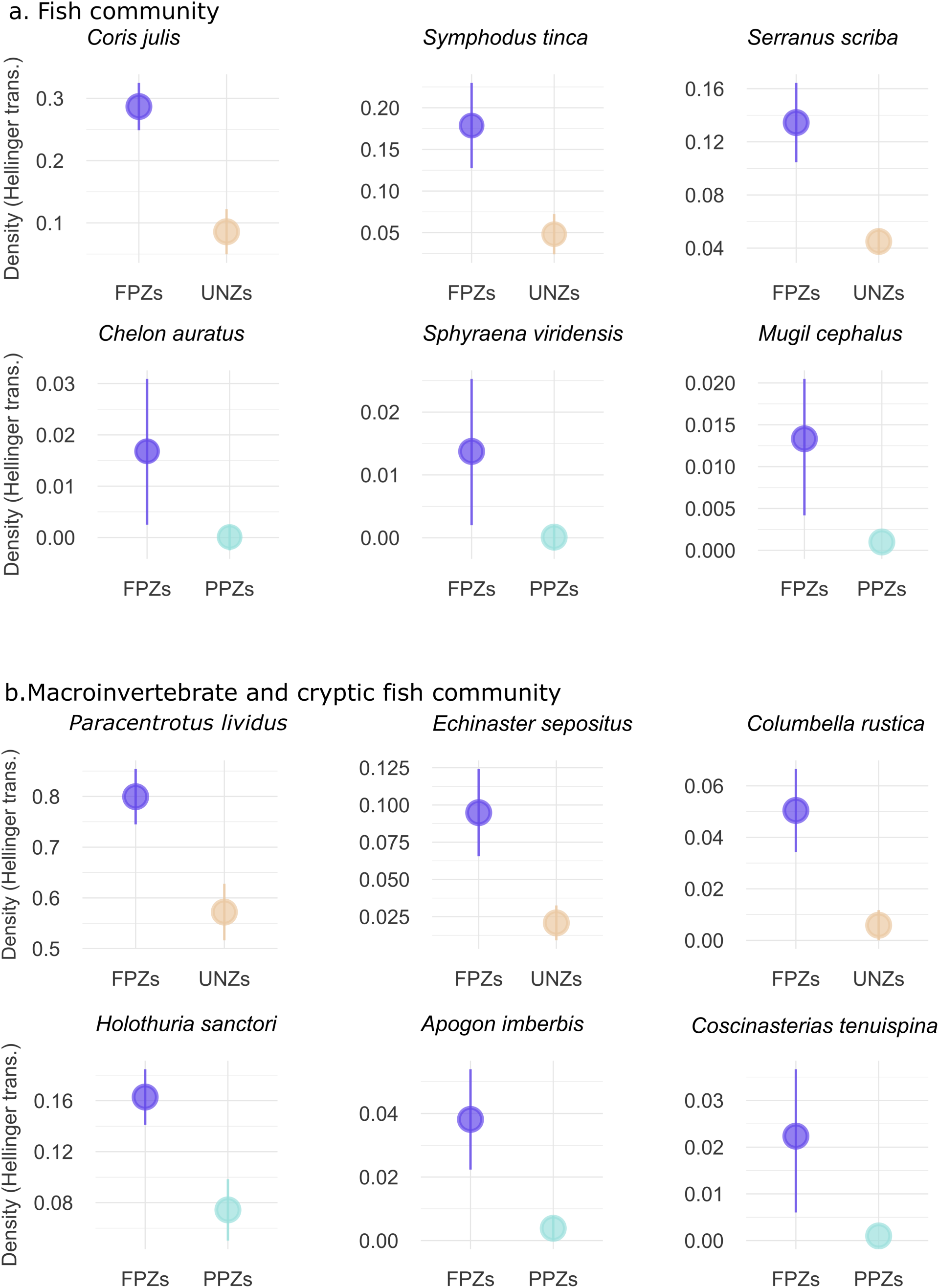
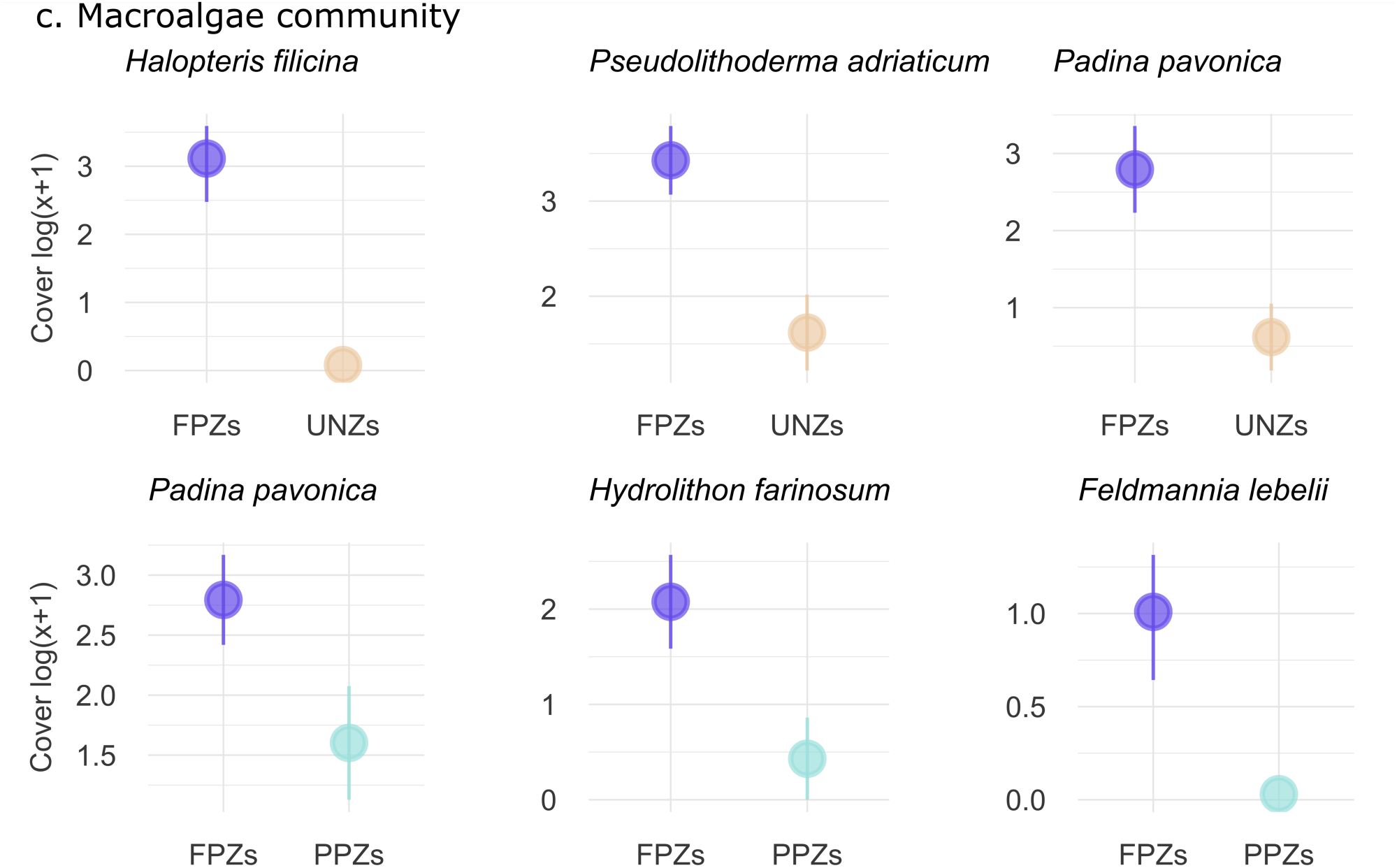
Representation of the three species with the highest density or coverage within the a) fish, b) macroinvertebrate and cryptic fish, and c) macroalgae community in the FPZs and their comparison between different protection levels. All these comparisons showed significant results p<0.05.

## 4. Discussion

Marine Protected Areas are an essential tool to safeguard marine biodiversity (Hilborn, 2016). Currently, we are entrusting the future of marine biodiversity to these marine protected areas, but there is still room for substantial improvement in their functioning (Edgar et al., 2014). In line with the necessity to improve the functioning of the marine protected areas, the metric presented here could help to monitor the protected areas based on vital ecosystem properties, such as resilience. In the present study, we showed that geographic position and environmental variables did not modulate the biological communities along the protection gradient. We also quantified the straightforward proxy measure of the ecological community recovery cost for fish, macroinvertebrates, cryptic fish, macroalgae, and the overall community, which is needed to move from unprotected or partially protected areas to fully protected areas. Besides, we unveiled which species increased their density or were only present in the areas of maximum protection. Most notably, our study provides a novel and readily quantifiable metric for considering the resilience of biological communities in conservation strategies. This approach also could be used in monitoring and designing terms in the field of conservation, because it allows to know the biological recovery cost before establishing the protected area, bringing resilience thinking to the decision-making table.

Currently, 8.2% of the ocean is under protection due to the existence of 18,444 marine protected areas (UNEP-WCMC, 2023). The actual protection of these areas varies depending on their restrictions and enforcement, among other factors. For example, there are zones with intermediate restrictions, called partial protection zones (PPZs in our study), where certain selective fishing gears and sports, such as scuba-diving, are allowed. There are also zones in which human activities are prohibited, called the no-take or fully protected zones (FPZs in our study). The latter are the areas that provide most benefits to marine biodiversity (Edgar et al., 2014; Sala and Giakoumi, 2017). Paradoxically, these no-takes zones are the least present in our seas, accounting for only 2.9% of the protected surface area (Marine Protected Atlas, 2023). Our findings show that the marine community varied with protection, i.e., between UNZs, PPZs, and FPZs. However, Sciberras et al. (2013) did not find differences in marine fish density between partially and fully protected areas. Conversely, fish biomass was higher in fully protected than in partially protected areas (Sciberras et al., 2013, Campbell et al., 2017). Also, the structure of the macroinvertebrate and cryptic fish and macroalgae community is well-differentiated between UNZs and FPZs (Shears and Babcock, 2003). Overall, FPZs was the protection figure with the largest biodiversity benefits, and it is a vital instrument for conservation (Costello and Ballantine, 2015; Sala and Giakoumi, 2018). Nevertheless, even today, selecting a geographic area to be protected remains a challenge. Selig et al. (2014) proposed a methodology for detecting priority areas to be preserved based on species richness and human pressure. Also, Edgar et al. (2014) revealed the five factors that make a marine reserve effective, so applying these factors could shed light on the detection of these priority areas to conserve. Unfortunately, the establishment of protection figures based on community key properties, such as resilience, remains scarcely studied and its application cannot be postponed.

Establishing conservation strategies based on biological properties targeting resilience, such as the ecological recovery cost, could increase the efficiency of protection figures (Côté and Darling, 2010). Our results showed that the ecological recovery cost demanded by the overall community (i.e., fish, macroinvertebrates and cryptic fish, and macroalgae) to move from partially protected to fully protected zones is less than the recovery cost required to go from unprotected to fully protected (0.32 *vs.* 0.42, respectively). Regarding the fish community, the ecological recovery cost for unprotected zones to resemble a community in a fully protected zone was lower than in the case of partially protected zones (0.12 *vs.* 0.23). A first plausible, but unlikely, scenario is that ecological recovery cost reaches low values because the unprotected zones had a biological composition equivalent to the fully protected zones. A second and perhaps more likely scenario might be the occurrence of illegal fishing activities in the fully protected zones (Edgar et al., 2014). This fact would explain why the ecological recovery cost between unprotected and fully protected zones is lower, despite significant differences between communities. A third scenario might suggest that partially protected zones are subject to significant pressure, which moves them away from the fully protected zones (since certain extractive human activities are permitted), driving the ecological recovery cost higher than that of the unprotected zones. This fact is undoubtedly of concern and must be addressed. Due to their high mobility capacity, fish may mask the clearer and robust trends found in macroinvertebrates and cryptic fish or macroalgae (Sanabria-Fernandez et al., 2018). A potential way to clarify what is happening with the fish recovery cost is to study illegal fishing pressure within the fully protected zones (Arias et al., 2015; Harasti et al., 2019). Moreover, establishing recurrent marine biodiversity monitoring programs could provide extremely robust and enlightening results (Bates et al., 2013).

Unlike the fish community, macroinvertebrate and cryptic fish feature reduced mobility (generally speaking) and may reflect robust spatial patterns (González-Duarte et al., 2014; Sanabria-Fernandez et al., 2018). In this sense, the recovery cost found in partially protected communities to resemble fully protected communities was 0.09, the smallest value found in the study. The composition of both communities was similar and statistically not different. On the contrary, the recovery cost of unprotected zone communities to become fully protected zones communities was 0.18, supported by significant differences between both communities. A probable interpretation of this interesting result is the existence of non-extractive pressures on the macroinvertebrate and cryptic fish community that may cause the degradation of this benthic community. For example, aquaculture installations (Silvert, 1992; Hargrave, 2010) or desalination plants (Ruso et al., 2007) could modify the nearby benthic community. This result indicates that changes in the macroalgae community demand a high cost in biological terms. González-Duarte et al. (2014) showed that sessile communities are faithful indicators of the state of the environment. Therefore, our study may be showing signs of some non-extractive human pressure in this area, causing modifications on the entire macroinvertebrate and cryptic fish, and macroalgae communities.

In this research, we considered the ecological community recovery cost as a proxy of the multidimensional distance from a degraded community to the confidence interval of the centroid of a healthy community. In other words, the path one community should follow in order to resemble the reference one. We developed this approach from the standpoint of multivariate distances involving the entire biological community, i.e., considering species and their density, based on Shade et al. (2012). Concerning the distance measurements, Vasilakopoulos and Marshall (2015) and Vasilakopoulos et al. (2017) considered the concept of univariate distance to the tipping point as the community’s relative resilience to environmental stressors. In contrast, our metric goes beyond univariate distances and involves multidimensional measurements, providing a comprehensive dimension of its resilience. Besides using distances to shed light on resilience measurement, some other approaches integrate the anthropogenic, environmental, and biological dimensions to quantify it and unveil priority areas to preserve. For example, Green et al. (2009), Davies et al. (2016); Sanabria-Fernandez et al. (2019) integrate the human, environmental, and biological dimensions in the detection of highly resilient marine areas in tropical and temperate latitudes. However, these studies pose a global resilience perspective of the system encompassing multiple factors and possibly masking an important fraction of the biological reality. On the contrary, our approach manages to obtain a faithful and clear vision of the cost of biodiversity recovery using community datasets. Specifically, it considers the entire biological composition of the community and involves the density of each species without distorting its reality. Also, there are some caveats associated with the methodology presented here, which might limit the generalizability of the findings. I) our results include an inherently non-significant fraction of geographic and environmental effects on the dissimilarity measures (see section 3.1). However, it is common that multiple environmental and geographic factors could also be contributing to this dissimilarity quantification between protected biological communities distributed across space. Thus, taking out its effects should be a preliminary step in these cases. II) In terms of temporal scope, this research is a temporal snapshot. However, the lack of long-term time series quantifying marine communities makes this potential limitation a strength by providing a benchmark for upcoming empirical studies seeking to understand the ecological recovery cost of marine communities over time. III) The proposed resilience measurement metric is founded on linear concepts; however, it is important to highlight that our current efforts are directed toward the development of nonlinear approaches to the study of the resilience of the communities.

The metric introduced in this study has direct and potential applications in biodiversity conservation in marine and terrestrial ecosystems. For example, this tool can help detect areas with low values of ecological recovery costs, which are strong candidates for protection. This fact implies redefining conservation policies, since now conservation agencies and policy-makers are able to perform preliminary estimates of the ecological community recovery cost. It enables us to elaborate on different recovery strategies, depending on the conservation aim and ecological recovery cost of each area. To satisfactorily apply this approach, a broad and intense study of the geographical area is necessary, but far from being a drawback, this would imply a benefit for unprotected areas, since it requires implementing biodiversity monitoring and assessment programs. Besides, the metric suggested here has a wide range of applications in terrestrial, freshwater, and marine ecosystems at spatial or temporal scales, depending on the study’s goal. Overall, our approach poses a robust tool that could help prioritize areas prone to improvement in biodiversity based on their resilience, contributing to more efficient conservation measures.

## Acknowledgments

We are honestly grateful for the direction Parque Natural Cabo de Gata-Níjar for their willingness to efficiently issue the scuba diving permits. JGA was supported by a Ramón y Cajal fellowship of the Spanish Ministry of Economy, Industry and Competitiveness (RYC-2016-20528). This article was written and submitted during JAS-F’s Margarita Salas postdoctoral research fellowship, funded by Universitat de Barcelona and the Ministerio de Universidades, and the Unión Europea Next generation, together with the “Plan de recuperación, transformación y resiliencia“. Additionally, we would like to express our appreciation for the insightful comments provided by Dr. Natali Lazzari which greatly enhanced the quality of this study.

---

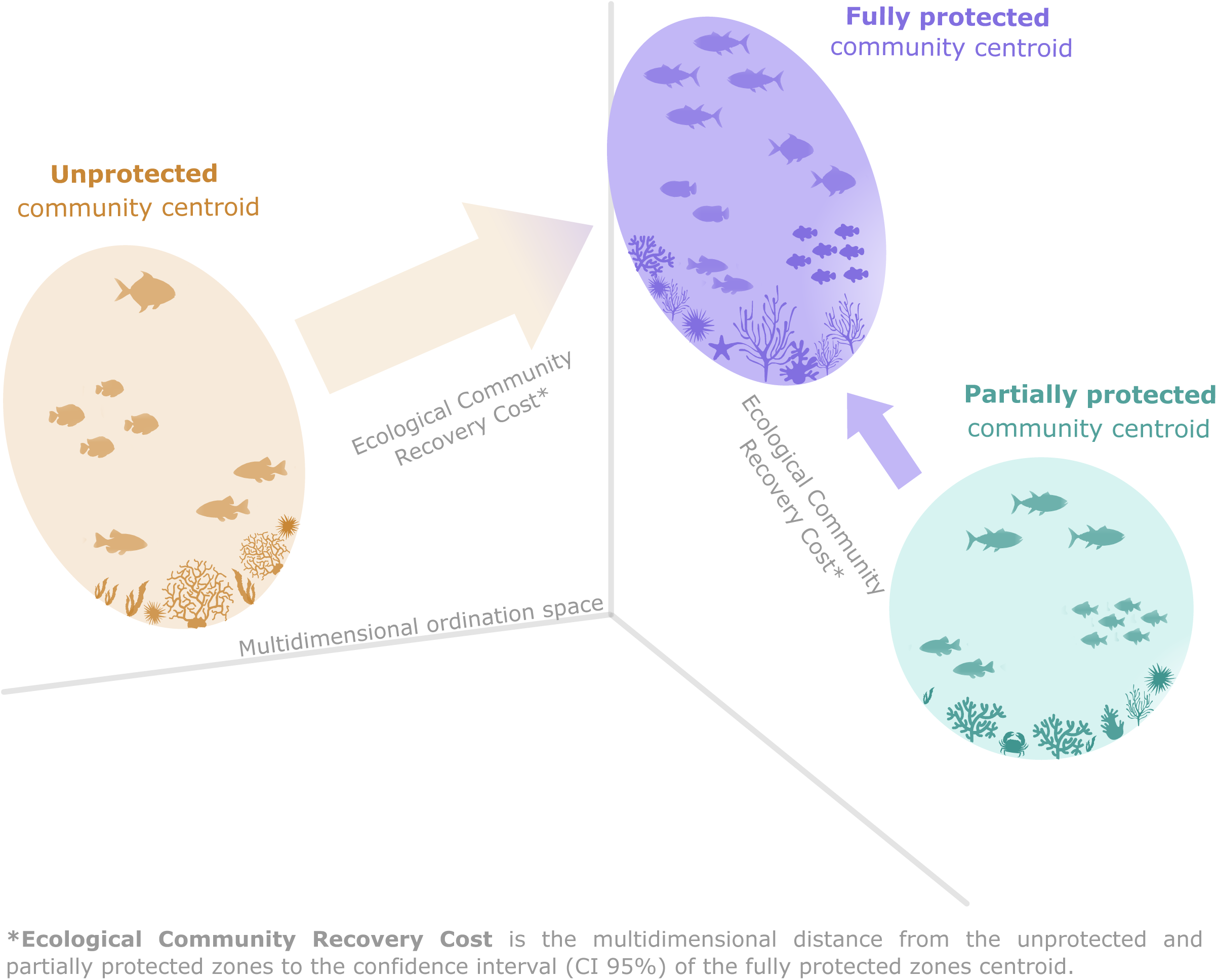

## Notes

https://www.reeflifesurvey.com

### Competing Interest Statement

The authors have declared no competing interest.

