## Supplementary material for "Marine protection enhances the resilience of biological communities on temperate rocky reefs"

**Supplementary Table 1**. Summarized results of Mantel test performed between geographic position and biological communities. We also ran the Mantel partial test, including the abiotic variables and biological communities. Here we controlled the effect of the geographic position by adding the geographic position of the locations. The $r_M$ is the Mantel statistic r.

| | Mantel test | | Mantel partial test | |
|---|---|---|---|---|
| **Biological community** | **$r_M$** | **P-value** | **$r_M$** | **P-value** |
| Overall | 0.1 | 0.06 | -0.13 | 0.954 |
| Fish | 0.03 | 0.21 | -0.04 | 0.69 |
| Macroinvertebrates and cryptic fish | 0.04 | 0.2 | -0.08 | 0.92 |
| Macroalgae | 0.11 | 0.06 | -0.02 | 0.601 |

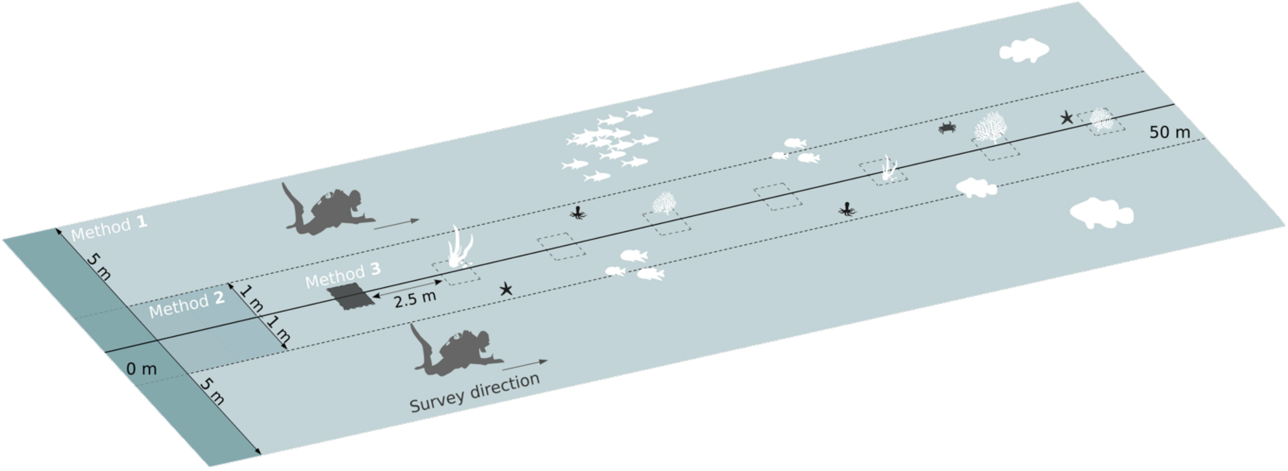


**Supplementary Figure 1.** Reef Life Survey protocol outline. Representation of the three approaches used and their technical specifications.
